# Unravelling the relationship between isolated bone matrix vesicles and forming mineral at the nanometer scale

**DOI:** 10.1101/2023.05.09.539570

**Authors:** Marcos A. E. Cruz, Luco Rutten, Martijn Martens, Onno Arntz, Fons van de Loo, Elena Macías-Sánchez, Anat Akiva, Pietro Ciancaglini, Ana P. Ramos, Nico Sommerdijk

## Abstract

Matrix vesicles (MVs) are involved in the initial deposition of hydroxyapatite (HAp) during bone mineralization, but their mechanism of action is not yet fully understood. *In vitro* studies propose two pathways by which MVs can trigger HAp precipitation: the first is mediated by their enhanced phosphatase activity, and the second suggested to depend on structural components present in MVs to mediate nucleation directly from soluble ions without the requirement of phosphatase activity. However, the relevance of these two pathways for bone mineralization and the relationship between MVs and forming mineral in such *in vitro* experiments remains unclear. Here, we used near-native cryoTEM nanoscale imaging in combination with bulk characterizations to disentangle the content and action of MVs during *in vitro* mineralization. We show that MVs isolation by conventional ultracentrifugation results in heterogeneous dispersions containing non-vesicular particles, including collagens and proteoglycans, in addition to bilayered vesicles. The separation of phosphatase-enriched MVs from non-vesicular particles and comparative mineralization experiments demonstrated that the ability of MVs to induce fast mineralization, independently of phosphatase activity, depends on the presence of non-vesicular particles. Therefore, we conclude that the primary pathway by which MVs trigger mineralization is through the action of their resident phosphatase enzymes, with the direct mineral nucleation to be a secondary event consequential of their membrane components. Lastly, we observed mineral formation restricted to the extravesicular space or in close proximity to the membrane interface, suggesting that the relationship between MVs and forming mineral is more intricate than previously understood.

## 1. Introduction

Bone biomineralization is the process by which calcium phosphate in the form of carbonated hydroxyapatite (HAp) is deposited in and on collagen fibrils within the extracellular matrix^1^. Despite its significance, the mechanisms behind the initiation of HAp mineralization within the collagen matrix are still not fully understood. MVs were first imaged in the late 1960s^2, 3^, when examining cartilage mineralization during endochondral bone formation. At the ultrastructural level, MVs are membrane-enclosed 100-300 nm nanostructures containing mineral associated to the lipid bilayer both internally and externally, always identified where HAp is initially deposited within the ECM^4^. However, the precise visualization of MVs within tissues depends on the methodology applied for sample preparation and imaging^5^. Furthermore, 2D electron microscopy images of tissue sections are limited in providing accurate 3D positioning between vesicles and minerals^6^. Despite the controversies imposed by these technical limitations, MVs have been extensively demonstrated to be present in different tissues undergoing mineralization, such as calvarial osteoid^7^ and dentin^8^. They have also been observed in pathological mineralization^9^, and it is now widely recognized that MVs play a role in the initiation of mineralization. However, the mechanism of action remains largely unknown.

The controversy surrounding the function of MVs is partly due to the difficulties in investigating them in detail. Their size in the nanometer scale and their presence in the densely packed ECM make reliable characterization challenging. Therefore, the main approach to study MVs is by isolating them from native tissues and reconstituting their content and functionality (i.e. mineralization) *in vitro*. A seminal approach used enzymatic digestion of epiphyseal cartilage, along with differential ultracentrifugation steps, to extract a vesicle pellet containing MVs^10^. Application of this approach revealed one of the main features of MVs, that is their increased alkaline phosphatase activity when compared to the mother cells^10^. Optimizations of this methodology ultimately succeeded in isolating a MVs fraction that was both reproducible and capable of inducing fast mineralization in solution^11^. This methodology is now the gold-standard to study the *in vitro* functionality of MVs and has been extensively applied for chicken and mouse embryonic bones.

The ultimate goal of isolating MVs from tissues is to retrieve their ability to induce mineralization *in vitro*. Inspired by their enrichment in alkaline phosphatase and adenosine triphosphatase (ATPase) activity, these experiments started by exposing MVs to a medium containing organophosphate compounds, such as ATP^12^. The ability to degrade organophosphates compounds has for long being implicated in the bone formation^13^. This enzymatic control has a dual effect, not only removing the inhibitory effect of the pyrophosphate (PPi), that strongly interacts with Ca^2+^ and supresses mineral formation and growth^14^, but also by the ability to produce free phosphate ion (Pi) to trigger mineralization. Exposure of MVs to a medium containing organophosphates induced the precipitation of calcium phosphate mineral^15, 16^. It is now recognized that the enzyme tissue non-specific alkaline phosphatase (TNAP) responds as the more important phosphatase operating in MVs. Enzymatic cleavage of TNAP from the outer membrane of MVs resulted in 80% reduction in mineral deposition^17^. The action of TNAP evidences a major role of MVs by fine-tunning the promotor/inhibitor molar ratio, Pi/PPi, permissive for mineralization^1, 18^. Failure in the control of inhibitors’ concentration is related to the development of several mineralization disorders and skeletal abnormalities^19^. Moreover, in cooperation with TNAP, other phosphatases have also been demonstrated to be operational within MVs, such as the nucleoside pyrophosphohydrolase-1 (NPP1)^20^ and the orphan phosphatase (PHOSPHO1)^21^. Alterations in the function of these phosphatases lead to softening of bone, spontaneous fractures, loss of teeth, as well as pathological calcification of soft tissues^18, 20, 22^. Remarkably, MVs isolated from wild-type and TNAP-, NPP1-and PHOSPHO1-knockout animals showed different ability to degrade organophosphate substrates and thereby to trigger *in vitro* mineralization^23^.

In spite of their intrinsic phosphatase activity, MVs have also been demonstrated to induce mineral precipitation *in vitro* without the need of any organophosphate degradation^24^. In this case, mineralization is described to occur by simply uptaking Ca^2+^ and Pi ions from the medium into the newly formed mineral through a nucleation process driven by MVs structural components. These components, referred to as a “nucleational core” are hypothesized to be composed of a membranous component formed between phosphatidylserine and annexins^25, 26^, and a pool of bound Ca^2+^ and Pi ions^27^. Isolated MVs are described to contain a large amount of Ca^2+^ and Pi, with the majority of the Ca^2+^ (>90%) initially present in an bound form, but only about 8-10% of the Ca^2+^ complexed with acidic phospholipids, as revealed by biphasic solvent partition of electrolytes^28^. This pool of bound Ca^2+^ and Pi ions was demonstrated to be a primordial component for the ability of MVs to trigger mineralization in a phosphatase-independent manner, since their removal upon treatments with slightly acidic buffer, calcium chelating agents or through sucrose fractionation hindered the functionality^29, 30^.

Historically, *in vitro* mineralization of MVs were termed “uptake” assays, in which the amount of Ca^2+^ and Pi ions incorporated into the newly precipitated mineral over time in the presence of the vesicles were radiometrically measured. These experiments aided to demonstrate the rate, type, and amount of mineral formed *in vitro*, as extensively reviewed by Wuthier^31^. However, a major challenge is the translation from bulk observations of *in vitro* mineralization to the underlying mechanisms by which enzymes and proteins operate within MVs, which occurs at a nanometric scale. Moreover, the relation between vesicles and forming mineral in such experiments is yet poorly understood. So far, visualization of mineral associated with MVs have been obtained with aid of conventional electron microscopy, but very few reports combined bulk assessments of mineral formation with imaging. Moreover, it is required to sediment MVs and mineral prior to the deposition of the collected material in TEM grids. This approach has allowed to follow the formation of mineral particles during in vitro mineralization. However, conventional TEM have the disadvantage of not faithfully preserve the imaged material in a fully hydrated state. This is of particular importance when handling biological samples like MVs, that include membranes and mineral particles, to prevent the collapse of the three-dimensional volume and extraction or mixture of the contents upon dehydration.

Here, in order to overcome these challenges and to solve in depth the relation between MVs and forming minerals, we used cryogenic transmission electron microscopy (cryoTEM) to image vesicles and mineral in a near-native state. CryoTEM involves the investigation of a thin vitrified film of a solution containing the particle of interest. During vitrification, all processes happening at bulk level are arrested and the particles under investigation become embedded in an amorphous film of the solvent^32^. Therefore, this imaging approach brings an unprecedent advantage of visualizing processes in a near-native state by keeping the whole material in a fully hydrated state. Application of cryoTEM was fundamental to uncover different aspects of bone biomineralization, such as the mechanism of collagen mineralization^33, 34^ and the phase transitions observed during HAp formation^35^. Therefore, cryoTEM can be explored as a powerful approach to investigate the content and functionality of MVs *in vitro*.

## 2. Results

### 2.1. Characterization of crude MVs isolated by differential ultracentrifugation

After enzymatic digestion of epiphysial cartilage slices with collagenase to release extracellular matrix-trapped vesicles, MVs were isolated using a standard three-step differential ultracentrifugation protocol (Figure 1a). Crude MVs (the pellet obtained at 80 000 *x g*) display all the characteristic features described in the literature^11^. A population of small vesicles with a diameter in the range of 100 – 300 nm was identified by nanoparticle tracking analysis (NTA) (Figure 1b), displaying high alkaline phosphatase (TNAP) activity (Figure 1c) and a typical electrophoretic protein profile (Figure 1d). From the SDS-page electrophoresis, we can discern 4 intense bands at apparent molecular weight of ∼ 30, 39, 42 and 45 kDa. The presence of these intense bands within this molecular weight range are characteristic of MVs isolated from chicken bones^11, 31^. The 30-and 33-kDa bands are now known to be annexin A5 (AnxA5); the 36-kDa protein is annexin A2 (AnxA2); and the 68-kDa protein is annexin A6 (AnxA6), but usually yielding a more faint band^31^. The presence of these proteins within chicken-derived MVs was confirmed by proteomic analysis^36^. TNAP appear as a faint and broad band at ∼ 70-76 kDa^24^, as also confirmed by western blotting analysis (Figure 1c). Other less intense bands appear migrating at higher apparent molecular weight (∼ 100, 150 kDa), previously assigned to collagen from the extracellular matrix^37^. Finally, cryoTEM imaging (Figure 1e) allowed visualization of the crude MVs preparations in a near-native state, indeed revealing a population of bilayered vesicles, with diameters in the range of 100-200 nm, corroborating NTA results. However, alongside the vesicles, a large amount of non-vesicular material was observed, most likely extracellular matrix components, including collagen fibrils (Figure 1f), and mineral particles with needle-like morphology typical for calcium phosphates. These mineral particles were either associated to the vesicles (green arrows, Figure 1e), or in aggregates associate to proteinaceous materials (Figure 1g). The nature of these mineral particles is uncertain. They might come from either mineralization triggered by MVs during the digestion step or endogenous mineral particles.

**Figure 1.**
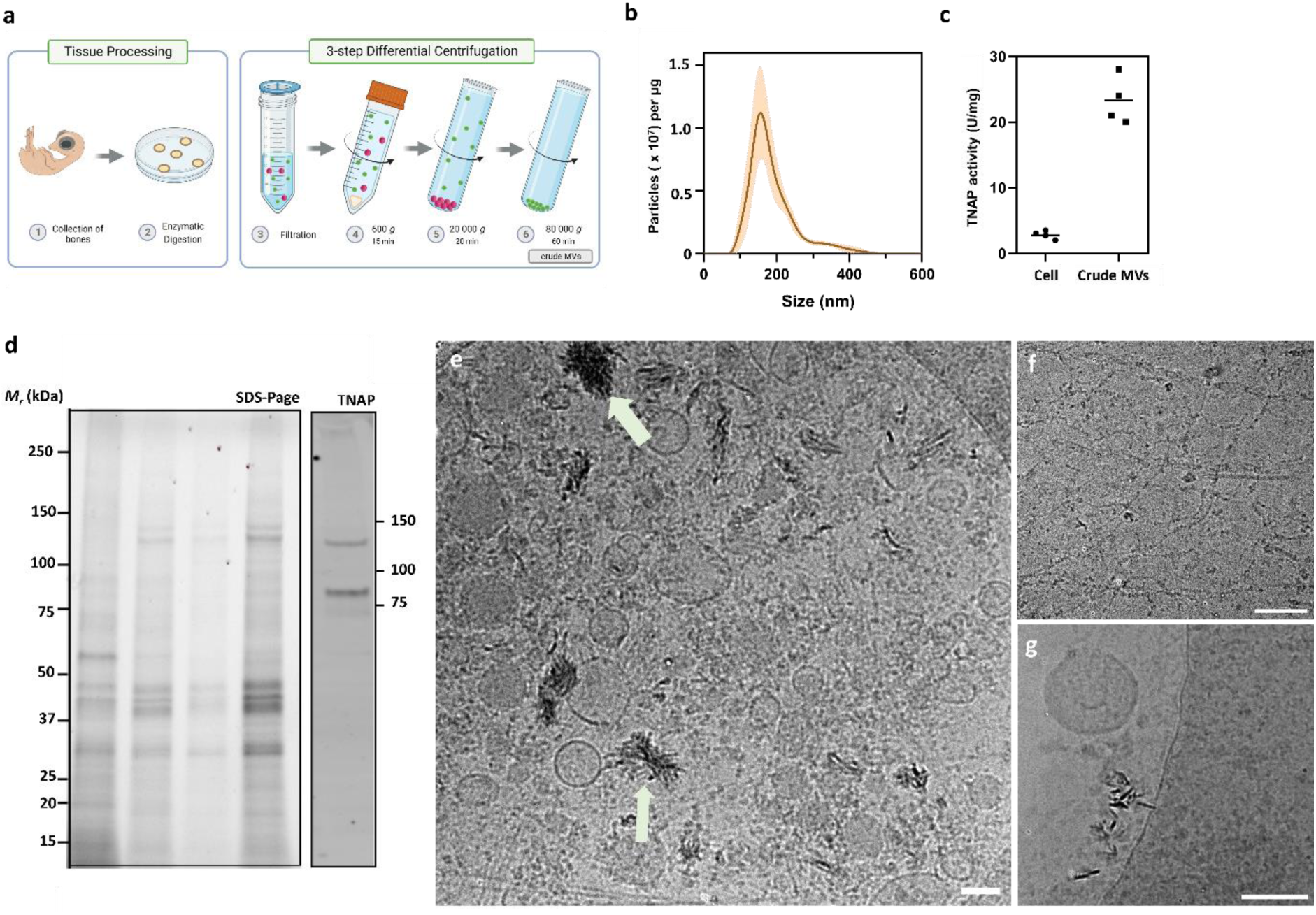
Characterization of crude MVs preparations. a) Crude MVs were isolated after enzymatically digestion of bone slices and collected through differential ultracentrifugation. b) Distribution of particles size measured by NTA. Data presented as number of particles per µg of total protein, obtained for 3 individual experiments. c) TNAP activity (U/mg) of crude MVs compared to cell pellet (step 4 in the scheme presented in the panel a). d) SDS-PAGE migration pattern of crude MVs for 4 independent experiments (3 µg of protein per lane). The overall migration profile observed for MVs is in accordance with the expected from the literature. Notice the reproducibility in the protein profile observed for 4 different crude MVs preparations. Western Blotting against alkaline phosphatase (TNAP) revealed the presence of a broad band migrating ∼ 75 kDa, and a band at ∼ 130 kDa. The broad band is related to the high glycosylation of TNAP, while the band at ∼130 kDa is assigned to the dimeric form of the enzyme, that is the active conformation^38^. e) cryoTEM imaging reveal the heterogeneity of crude MVs. A mixture of bilayered vesicles with non-vesicular particles is clearly discernible. f) non-vesicular particles are dominated by fibrillar proteins. While some mineral particles appear to be associated to vesicles (white arrow, panel e), mineral particles are also observed associated to non-vesicular material (panel g). Scale bars, 100 nm.

### 2.2. Density-gradient fractionation separates MVs from non-vesicular particles

To further separate bilayered vesicles from non-vesicular components, we used a density gradient centrifugation. Crude MVs were bottom-loaded into an iodixanol density-gradient column (10-45%), to separate MVs from co-sedimented non-vesicular particles (Figure 2a). Iodixanol was chosen for being isosmotic in all density ranges used. Moreover, bottom-loading is advantageous since membranous vesicles will float upwards, leaving denser components downwards. After centrifugation at 120 000 *x g* for 16 h to assure the isopycnic separation, low-dense fractions were characterized by high TNAP activity (Figure 2b) and high number of particles detected with NTA per µg of protein (Figure 2c), when compared to crude MVs. High number of particles per µg of protein is an indicative of vesicleś purity, since the presence of high amounts non-vesicular associated proteins will decrease the number of particles normalized per protein amount^39^. However, high density fractions were characterized by low TNAP activity and reduced number of particles/µg of protein. Therefore, low-dense MVs with high TNAP activity represents functional vesicles implicated in bone biomineralization. SDS-PAGE comparison of the protein profile of crude and low-dense MVs revealed remarkable differences (Figure 2d). Low-dense MVs lack bands at high molecular weight range (100-150 kDa) and some major bands at 38 and 46 kDa compared to crude MVs. Previous studies described that treatment of crude MVs with hypertonic salt solutions selectively removed bands on SDS-PAGE analysis at an apparent molecular weight of 130-150 kDa, identified to be related to type II collagen and its fragments, as confirmed by antibody recognition^37, 40^. Further extraction with low ionic strength solutions removed two major bands on SDS-PAGE analysis of MVs, at an apparent molecular weight of 40-48 kDa, identified as proteoglycan-related proteins^40^. The main changes in the SDS-PAGE profile of low dense MVs were found close to these apparent molecular weight range, thus suggesting that collagens, proteoglycans, and associated proteins correspond for the majority of non-vesicular particles present in crude MVs. Finally, we imaged low dense MVs with the aid of cryoTEM, confirming the presence of single-membrane bilayered vesicles with diameters in the range of 100-200 nm. CryoTEM images also confirmed that low dense MVs are devoid of any large macromolecular aggregates and fibrillar proteins, corroborating NTA and SDS-PAGE data. Moreover, we also observe that all mineral particles observed in crude MVs (Figure 2e) were removed after density fractionation, and no evidence of calcium phosphate mineral presence was find neither in the inner nor at outside of the vesicles.

**Figure 2.**
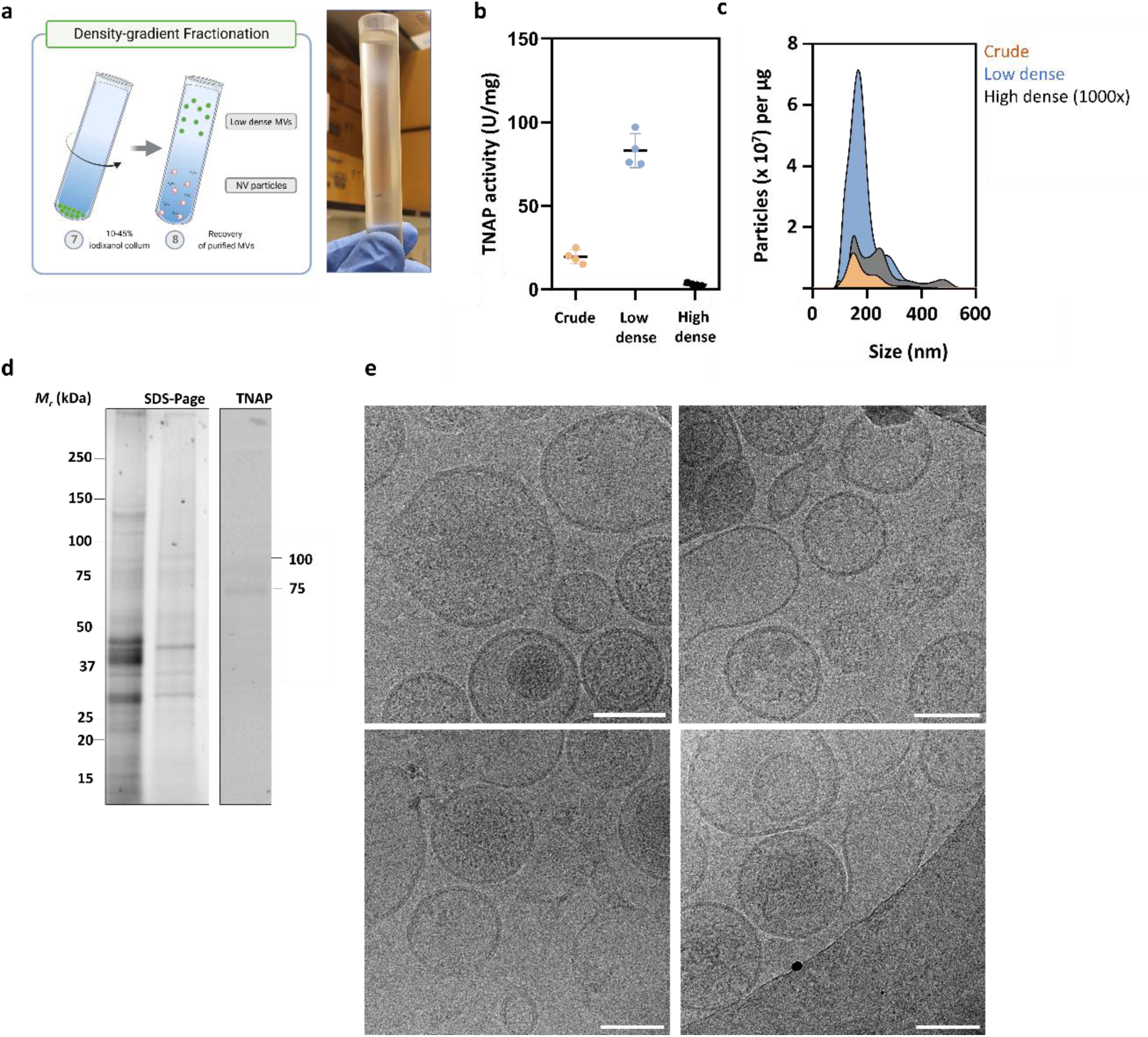
Iodixanol density-gradient was able to separate low dense MVs from non-vesicular components. a) Crude MVs were bottom-loaded in a 10-45% iodixanol column, that after centrifugation for 16 h separated low-dense MVs from non-vesicular components. Photography of the gradient column after the centrifugation showing a broad band containing the low-dense MVs at the interface of 20-22% solutions. b) TNAP activity (U/mg) for low dense MVs compared to crude MVs. d) SDS-Page electrophoresis gel comparing crude MVs with low dense MVs. e) cryoTEM images of low dense MVs showing a distribution of bilayered MVs with almost absence of non-vesicular particles. Notice that the morphology of vesicles are heterogenous, with some vesicles appearing electron-lucent, others electron-dense, and some minor extent of multi-layered, that might be associate to the fusion upon ultracentrifugation^41^. Scale bars, 100 nm.

### 2.3. Crude and low-dense MVs show different kinetics of induction of mineralization from soluble Ca^2+^ and Pi ions

Crude and low-dense MVs were exposed to SCL containing Ca^2+^ and Pi ions, reproducing conditions where fast induction of mineralization without the requirement of phosphatase activity is described to occur^11, 24^. Then, we used cryoTEM to examine mineral formation in the presence of both the MVs preparations. To minimize bias from disturbance of the samples, we avoided the need of centrifugation to collect MVs and minerals at specific time points by conducting an on-grid mineralization experiment. In this approach we incubated MVs in solutions dropped directly on a TEM grid, in a humidity-controlled environment, and further processed for vitrification at a desired time point.

After 24 h, cryoTEM images confirmed the formation of crystalline mineral in crude MVs exposed to the SCL medium containing Ca^2+^ and Pi ions (Figure 3a-b). The crystals appeared as large aggregates around the vesicles, and many intact vesicles that had no sign of any associated mineral were also observed (Figure 3b). In contrast, for low-dense MVs at same conditions, after 24 h, we observed the majority of vesicles devoid of any associated mineral (Figure 3c) and some of them associated to an amorphous-like mineral at their membraneś surface (Figure 3d). Note the presence of calcium phosphate clusters with approximately 20 nm, typical of metastable solutions still present in the samples containing low dense MVs even after 24 h (white arrows, Figure 3d). These observations were confirmed throughout the entire volume analysed, as demonstrated by the low magnification images in Figure 4.

**Figure 3.**
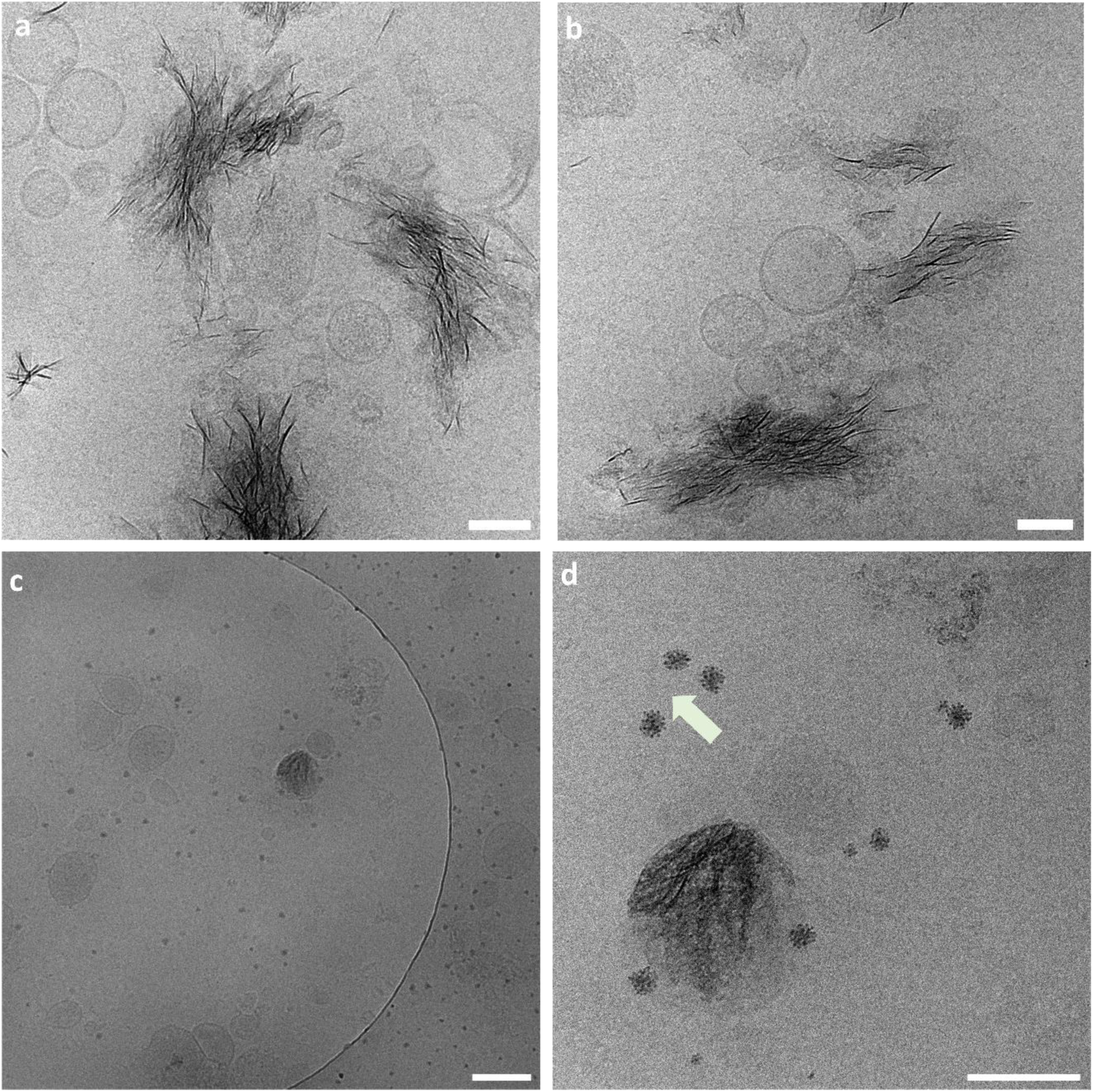
CryoTEM reveals the relationship between MVs and forming mineral. CryoTEM images of crude (a,b) and low-dense MVs (c,d) incubated in SCL containing Ca^2+^ and Pi ions for 24 h. Arrow in panel d points to calcium phosphate clusters. Scale bars, 100 nm.

**Figure 4.**
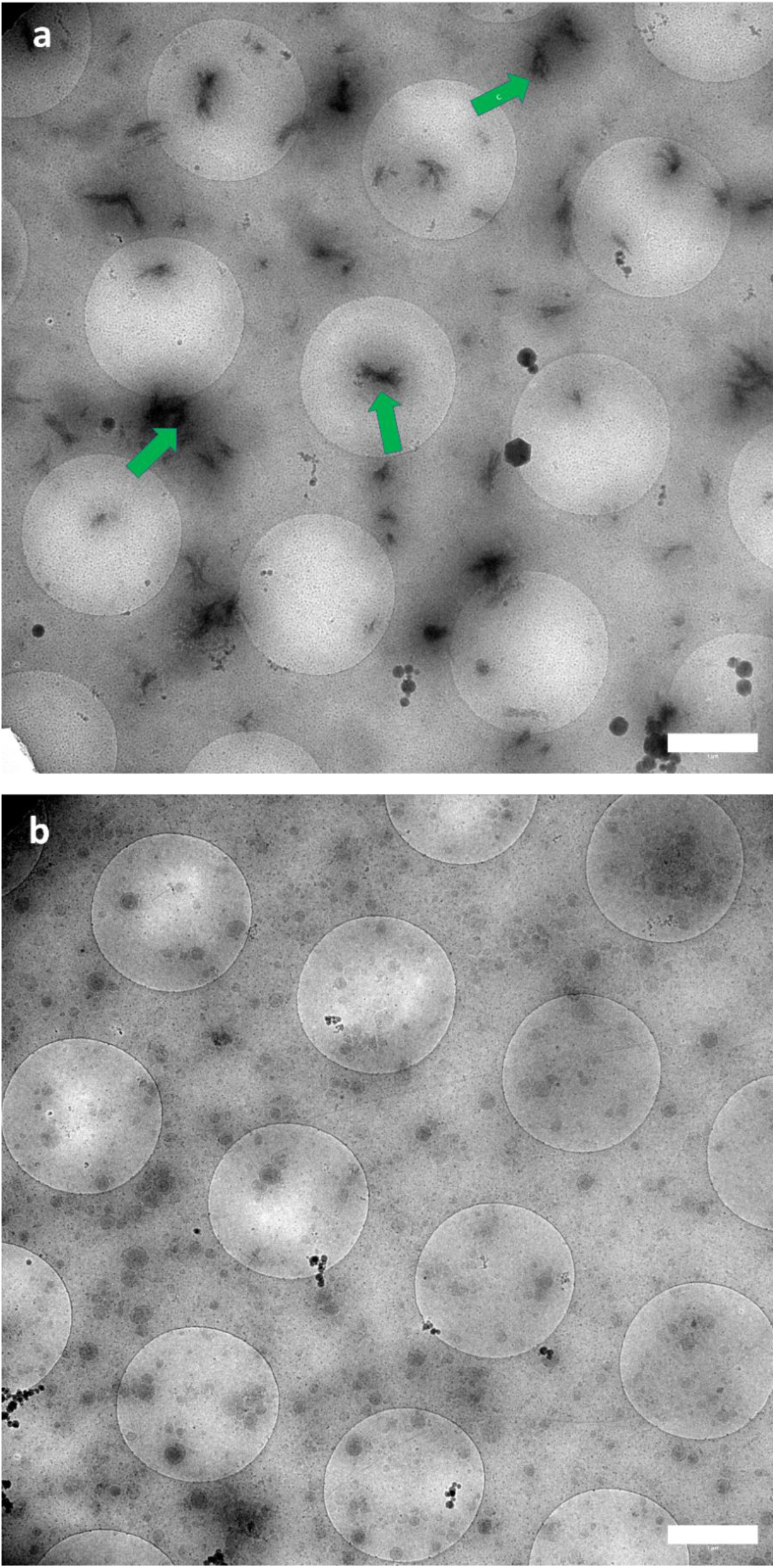
Low magnification images of crude (a) and low-dense MVs (b) induced for mineralization for 24 h. Green arrows in image a points the large mineral aggregates. Notice the remarkable difference in the number of vesicles, at same protein concentration, for crude and low dense MVs. Scale bar, 1 µm.

To improve the knowledge about the origin of the crystalline mineral present in the samples after 24 hours of mineralization, we conducted the experiments also in an earlier time point. We found that after only 8 hours of mineralization, crude MVs already showed evidence of crystalline mineral formation. These crystals were found to be associated with vesicles (Figure 5a) but were also dispersed freely in the medium along with intact vesicles that did not have associated mineral (Figure 5b). In a small fraction of the analysed sample, some vesicles were found to be intimately associated with mineral at their membranes (white arrow, Figure 5c). Similar results were observed for low-dense MVs, with vesicles without any signs of associated mineral dominating the sample (Figure 5d). Some amorphous-like mineral particles associated to the vesicleś membranes were also observed (Figure 5e), similar to the result found after 24 hours. Due to the high concentration of vesicles in low-dense MVs, we could often observe vesicles clustered together (Figure 5f).

**Figure 5.**
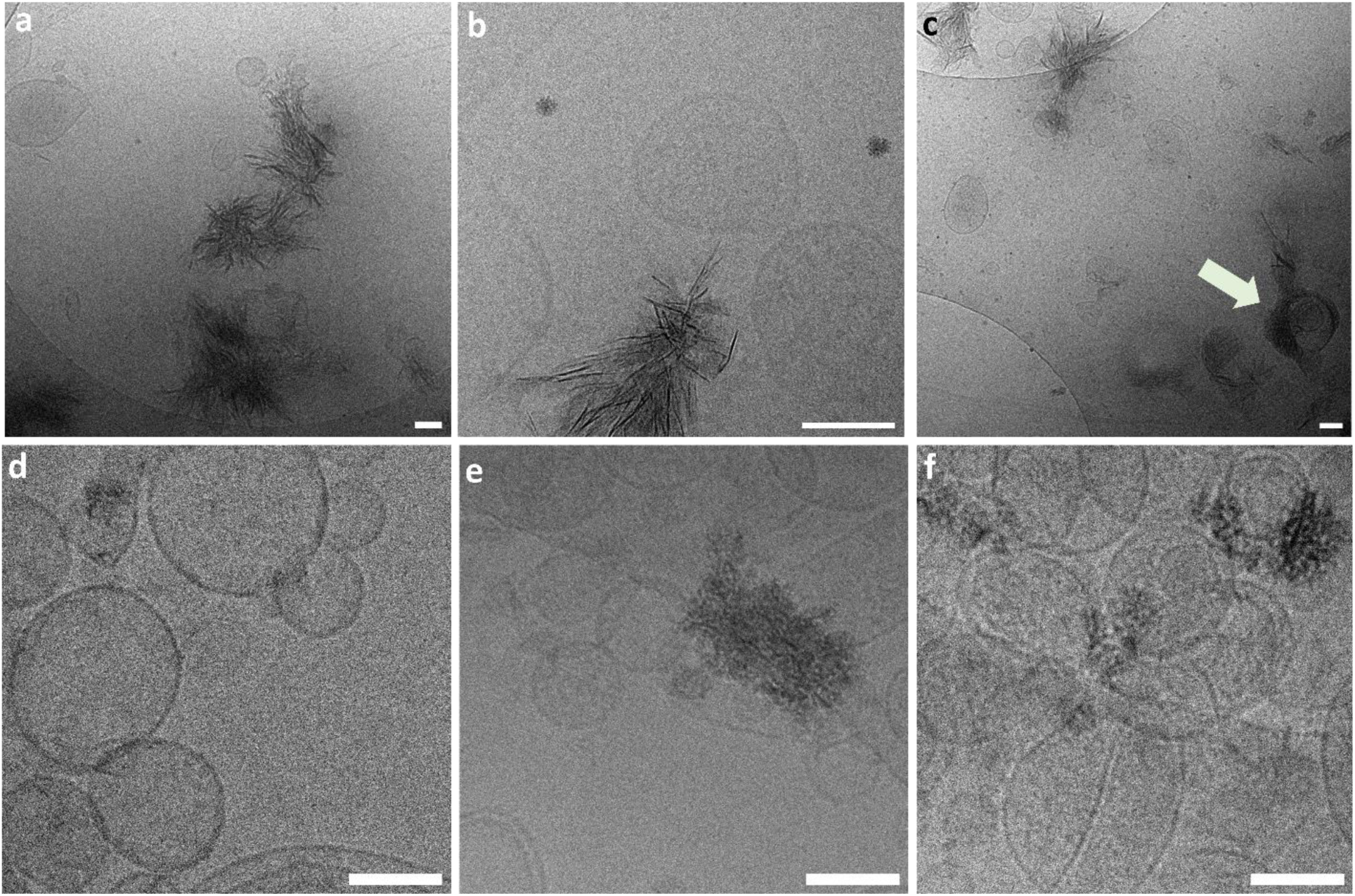
Relationship between MVs and forming mineral at early stage. CryoTEM images of crude (a-c) and low-dense (d-f) MVs incubated in SCL containing Ca^2+^ and Pi ions for 8 h. Arrow in panel d points to mineral associated to the membrane of vesicles. Scale bars, 100 nm.

To validate the findings by cryoTEM at the nanoscale, we also examined the differences in the ability of crude and low-dense MVs to trigger mineralization at the bulk scale. For this, we first used turbidimetry, since this approach has been extensively applied to probe mineralization of MVs *in vitro*. Tracking changes in the turbidimetry (absorbance at 350 nm) of the solution containing crude MVs (Figure 6a), we observed an initial increase in the turbidimetry after a lag time of 3 h, and an increase of 0.3 absorbance units after 8 h of incubation. However, low-dense MVs, at same protein concentration, induced slower changes in the turbidimetry of the solution, with an increase of only 0.1 absorbance units after 8 h of incubation. Turbidimetry is based on the measurement of the loss of intensity of transmitted light in a solution due to the scattering effect of particles in suspension. Therefore, an increase in the turbidimetry will be sensitive to changes in the diameter of scattering centres already present in the dispersion, or to the formation of new scattering particles in the medium. The turbidimetry measurements confirmed the cryoTEM findings, i.e., the formation of sub-micrometric clusters of particles in presence of crude MVs is responsible for the increase in the intensity of scattered light after incubation into SCL containing Ca^2+^ and Pi.

**Figure 6.**
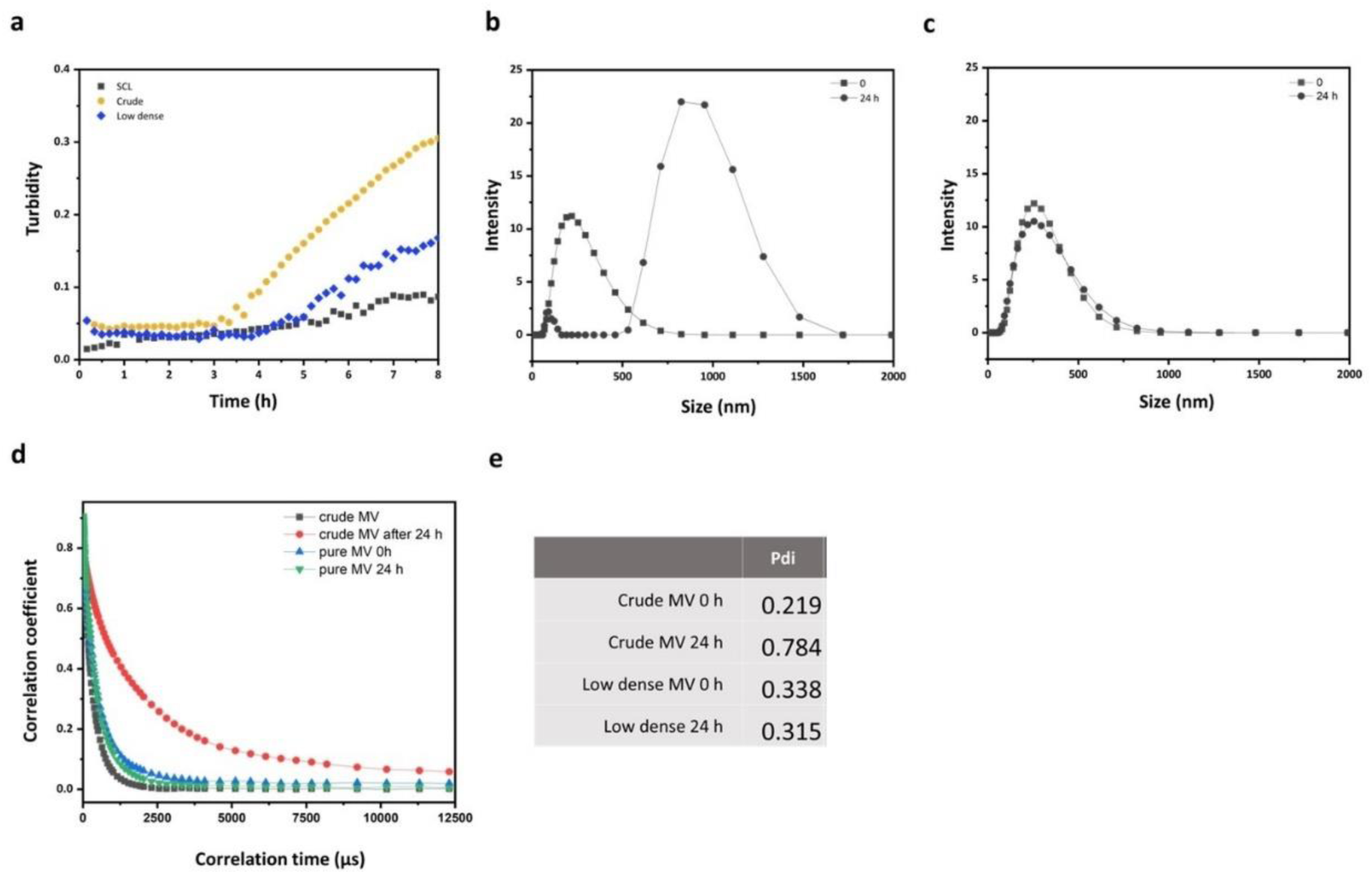
Examination of mineral formation at bulk level. a) Turbidity (absorbance at 350 nm) over time for crude and low-dense MVs in SCL containing Ca^2+^ and Pi, at 37°C. Distribution of particles measured with DLS at time 0, and after 24 h of mineralization, for crude (b) and low-dense MVs (c). d) Correlation coefficient versus correlation time (µs). e) Pdi values obtained for the different measurements.

To get a more quantitative data, we sought to use dynamic light scattering (DLS) to investigate the size distribution of particles in solution at the beginning of the mineralization experiment and after 24 hours of incubation. By DLS we measured the intensity of light scattered by particles dispersed in liquid. Fluctuations in the intensity of the scattered light as a function of time can be recorded and used to calculate de translation diffusion coefficient of the particles from a correlation function. Then, the diffusion coefficient is converted into hydrodynamic diameter of the particles by using the Stokes-Einstein equation^42^.

DLS data (Figure 6b) show that both crude and low-dense MVs exhibited one distribution of diameter centred at 250 nm in the initial time. However, after 24 hours of mineralization two distributions can be observed for crude MVs– one centred at 150 nm and another, much more intense, shifted towards particles with higher diameters. However, no significant changes were observed in low-dense MVs (Figure 6b), corroborating the cryoTEM observations, i.e., the formation of large mineral aggregates only for crude MVs after 24 hours of mineralization. Since the intensity of light scattering is proportional do the diameter^6^, bigger particles scatter more light than smaller ones, explaining why the distribution of particles was dominated by the bigger particles in the DLS intensity data for crude MVs after 24 hours. The differences observed in the size distributions become clearer when we analyse the raw correlation function (Figure 6d). The raw correlation data shows the correlation function plotted as a function of the correlation time. The exponential function reaches zero (loss of correlation) faster in samples containing smaller particles than larger particles^42^. Therefore, we can definitively conclude from the exponential curves in Figure 6d that crude MVs trigger the formation of larger particles after 24 hours of mineralization, while low-dense MVs remain almost unchanged corroborating at a bulk level our examination of mineral formation by cryoTEM. These observations were further confirmed by the polydispersity (Pdi) indexes obtained from the DLS data, which increased considerably for crude MVs samples after 24 hours due to the presence of large mineral particles (Figure 6e).

### 2.4. Crude and low-dense induce mineralization in presence of ATP

Finally, we tested the ability of the crude and low-dense MVs to induce mineral formation through by the hydrolysis of ATP by TNAP present in the MVś structure, to generate Pi. Since ATP is a strong mineralization inhibitor, no mineralization is induced without its degradation^43^. Both crude and low-dense MVs were able to hydrolyse ATP as revealed by the increased Pi concentration as a function of time (Figure 7a). However, the conversion rate is faster for low-dense MVs due to their increased TNAP activity compared to crude MVs. CryoTEM images of crude (Figure 7b-c) and low-dense MVs (Figure 7d-e) after 24 h of incubation in SCL in the presence of ATP revealed the formation of mineral precipitates around the vesicles, in accordance with previous reports^44, 45^.

**Figure 7.**
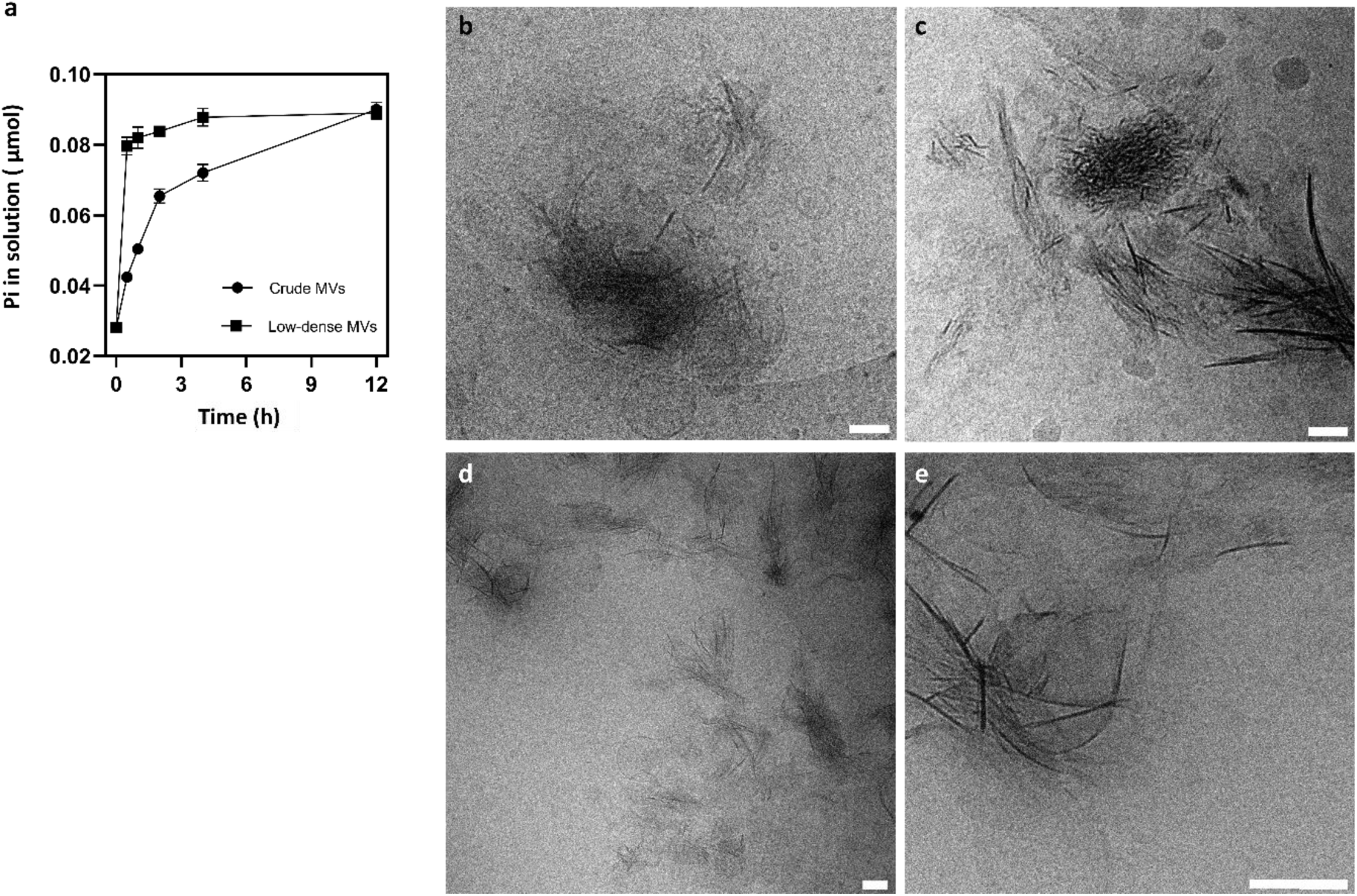
Mineralization induced by hydrolysis of ATP. a) Changes in the concentration of Pi formed as a function of time for crude and low-dense MVs exposed to SCL containing 1 mM ATP, at 37°C and measured in a reaction volume of 20 µL. CryoTEM images mineral formed when crude (b,c) and low-dense (d,e) MVs were incubated for 24 h in the SCL containing ATP. Scale bars, 100 nm.

## 3. Discussion

The coexistence of MVs along with apatite crystals within different tissues undergoing mineralization has been extensively demonstrated^31, 46^. However, understanding the mechanisms of MVs action is challenging due to the changes at nanoscale that takes place during bone mineralization. Furthermore, the formation of products resultant of MV action, i.e., HAp crystals, follows general physicochemical rules, thus requiring a multidisciplinary approach to fully appreciate all underlying mechanisms. The interpretation of the mineralization rate, type, and amount of mineral formed after exposure of MVs to mineralizing solutions has led to the proposal of two major pathways for HAp precipitation. These mechanisms involve either the direct nucleation from soluble ions, or through an increase in Pi concentration due to phosphatase enzymes able to hydrolyze organophosphate compounds. In this thesis studies, we presented new insights into these two mechanisms, thanks to our unprecedented capability to integrate bulk measurements with near-native cryoTEM imaging at a single vesicle scale.

First, we demonstrated that crude MVs preparations are heterogenous and contains a large amount of non-vesicular particles in addition to bilayered vesicles, based on cryoTEM images. Conventional TEM imaging of dried and stained vesicles revealed also the presence of fibrillar proteins in the samples but it was difficult to discern the relationship between the different components of the material (Supplementary Figure S1). By keeping the samples in hydrated state using cryoTEM, we observed that, in general, the fibrillar proteins are not intimately associated with the vesicles, but rather dispersed in the medium. Upon fractionation with the iodixanol-density gradient, we dissected a population of bilayered low-dense MVs absent of any non-vesicular structure. By comparing the mineralization ability of crude and low-dense MVs, we observed that removing these non-vesicular particles had a major impact in the onset of mineral formation in presence of soluble Ca^2+^ and Pi ions. Crude MVs induced fast formation of crystalline mineral after 8 h, however low-dense MVs promoted the formation of a small amount of mineral associated to their lipid membranes after 24 h, as observed by cryoTEM. This result is in accordance with previous studies in which fractionation of crude MVs in sucrose gradients yield a less-dense fraction of vesicles with high TNAP activity, but reduced ability to induce fast mineralization in the presence of Ca^2+^ and Pi ions^47, 48^. In those studies, it was demonstrated that Ca^2+^ and Pi ions migrated to denser sucrose fractions, not coexisting anymore with the TNAP-rich less-dense fractions. Since sucrose solutions are hyperosmotic at high concentrations, its use for MVs fractionation is thought to destabilize a luminal pool of Ca^2+^ and Pi ions due to the osmotic pressure imposed by high concentration of sucrose. Therefore, this pool of bound Ca^2+^ and Pi ions has been hypothesized to be a major component responsible for the ability of MVs to trigger mineral nucleation^27^. This hypothesis was supported by other treatments of crude MVs, such as citrate buffer, or with calcium-chelating agents, that also removed most of the associated mineral ions and as a consequence the ability of MVs to trigger mineral nucleation^27, 29, 49^. In our experiments we used a isosmotic medium (iodixanol) and added Ca^2+^ ions to prevent dissolution of labile ions^49^. Upon fractionation in the iodixanol gradient, analysis of cryoTEM images and the changes in the SDS-Page protein profile led us to the conclusion that this purification step removed most of the associated fibrillar collagens and proteoglycans from the samples. Therefore, our results suggest that the pool of bound Ca^2+^ and Pi ions, previously thought to be a luminal cargo of the bilayered MVs themselves, is rather associated to the non-vesicular particles (such as collagens and proteoglycans) present in crude MVs preparations. This reinterpretation also explains why the ability to induce fast mineralization could also be destroyed upon EDTA and citrate treatments, as we would not expect these highly charged molecules to cross the lipid bilayer to remove Ca^2+^ from the vesicle interior. Confirming our conclusions, Kirsch et al. reported that sucrose-fractionated low-dense MVs could recover the ability to induce fast mineral nucleation by adding native collagen to the medium^47^. Moreover, Wu et al. observed a marked reduction in the ability to trigger mineralization after treating MVs with hydrazine, that removed ∼95% of the proteoglycans associated with MVs^50^. Therefore, our observations suggest that non-vesicular proteins dominated by fibrillar collagens and proteoglycans responds for the ability of crude MVs to trigger mineral nucleation, either for bringing a large amount of Ca^2+^ and Pi ions, thus increasing supersaturation or through their intrinsic nucleating activity^51^. To support our conclusions, we confirmed that the fraction collected at higher density in the iodixanol gradient induced the formation of crystalline mineral after 24 h when exposed to the same conditions in which low dense MVs did not. (Supplementary Figure S2).

The ability of crude MVs to trigger mineral nucleation has been historically assigned to the existence of a nucleational core within MVs that is converted to apatite when in contact to Ca^2+^ and Pi ions. This nucleation core was hypothesized to contain two main components as follow: a pool of Ca^2+^ and Pi ions bound to luminal proteins and a membrane-associated complex of Ca^2+^, Pi, phosphatidylserine, and annexin that nucleates the formation of crystalline mineral^27^. Our results do not discard the possibility of intraluminal mineralization events during MV’s driven mineralization, however the association of mineral with the MV’s membrane confirm the importance of calcium-binding components to mediate mineral nucleation. The ability of Ca^2+^-binding sites to mediate mineral nucleation is a well-known phenomenon in biomineralization^52^. From cryoTEM images, we observed only a limited amount of mineral associated with the membranes of vesicles. While it is true that complexes of phosphatidylserine and annexins with Ca^2+^ and Pi can effectively nucleate HAp *in vitro*^25, 26, 53^, our results suggest that this capability might be of secondary importance when these components are present within the membrane of MVs, or occurring at a much slower rate.

The results also evidenced the primary functionality of MVs by making Pi available for mineralization through phosphatase activity. Upon exposure to ATP, both crude and low-dense MVs trigged mineral formation, as shown in the cryoTEM images, that revealed the formation of mineral crystals around the vesicles. TNAP played a significant role in this process, since it can efficiently hydrolyse organophosphate substrates and increase the concentration of Pi, which leads to the precipitation of minerals and the formation of minerals in the extravesicular space. The observation of minerals in the extravesicular space is in accordance with experiments conducted by Hsu & Anderson, in which MVs were exposed to an ATP-containing mineralizing medium for 24 hours to reach maximal mineral deposition, sedimented, incubated with a calcium-chelating agent (EGTA) for 24 hours to remove MV-associated minerals, and further sedimented to reveal vesicles that seemed to lack minerals but can still trigger precipitation upon ATP readdition^15^. Moreover, besides TNAP, MVs harbour other different phosphatases that share a common functionality: producing Pi for mineralization by degrading organophosphate compounds^21^. The origin of organophosphates found in the extracellular matrix is yet unclear. Some possibilities are related to polyphosphates^54, 55^, the cell-mediate secretion of ATP^56^, or the self-degradation of MVs’ membrane phospholipids^57^. These different phosphatases are demonstrated *in vivo* to work in non-reductant way, since either individual or mutual genetic ablation of their functions always result in a certain level of deficient extracellular mineralization. These observations together emphasize the primary function of MVs, which is to provide enzymatic machinery able to locally regulate the concentration of Pi. Mineral precipitation occurs when an optimized environment that allows mineralization is achieved, which involves supersaturation and consumption of inhibitors guided by principles of thermodynamics. At this point, a secondary role of MVs related to accumulation of Ca^2+^ can be also highlighted: Ca^2+^-binding sites of MVs act as motifs that interact with mineral ions and create boundaries through compartmentalization, changing energy barriers and creating smoother energy landscapes^58^.

Finally, another important insight from our observations is about the relation between MVs and forming mineral during *in vitro* mineralization. From ultrastructural studies of tissue sections, in which crystals were observed in and out the membrane of vesicles, MV-driven mineralization is hypothesized to follow a biphasic phenomenon, where mineralization is first initiated within the lumen of vesicles and then after membrane rupture, the crystals are released to extracellular medium^59^. Therefore, attempts have been made to relate the sequence of steps observed during *in vitro* mineralization to the *in vivo* mechanisms by which MVs initiate mineralization. Extensive data has been collected on the mineralization of MVs in the presence of Ca^2+^ and Pi ions, comprehensively reviewed by Wuthier^31^. By measuring the amount of mineral formed, it has been observed that MVs undergo a series of predictable stages. Initially, there is a lag period with minimal ion accumulation, followed by a period of rapid accumulation of Ca^2+^ and Pi. After this, there is an extended plateau period of slow ion accumulation, during which the Ca/P ratio gradually increases, approaching that of HAp after 24 hours. The first detectable mineral phase appears just after the period of rapid accumulation, which typically lasts 4 to 6 hours. This mineral phase is characterized by an acidic octacalcium phosphate (OCP)-like phase, with a Ca/P ratio close to 1.33, as determined by various spectroscopic methods^50, 60, 61^. The rationalization that the first detection of crystalline mineral after 4-6 hours reflects the biphasic process hypothesized to occur in MVs-driven mineralization has suggested a life span for these structures of short 4-6 h, after this membrane rupture should occur to grant the release the initially formed crystals. Herein, the cryoTEM images of crude MVs revealed that after 8 h, which from our turbidimetric analysis represents the end of the so-called rapid acquisition phase, crystals were mostly restricted to the extravesicular space, and intact vesicles could be detected even after 24 h of reaction. The lag period is currently believed to be a rate-limiting step, which reflects the inability of ions to pass through the vesicle membrane and reach the lumen. Once they reach the lumen, “sink conditions” provided by the nucleational core drive the accumulation of Ca^2+^ and Pi, that is converted into the first detectable OCP-like crystals. After careful combination of turbidimetry and cryoTEM experiments, we have reached the conclusion that the rapid acquisition phase observed does not accurately represent the spatial process by which Ca^2+^ and Pi ions are taken up into the lumen of MVs to trigger mineralization. Rather, it appears to be a bulk event of nucleation occurring within the solution, triggered by the nucleating activity of non-vesicular components present in crude MVs. Therefore, this reinterpretation of bulk mineralization experiments suggests that MVs have a significantly longer lifespan than previously thought.

SCL is a metastable solution stabilized against precipitation in the absence of a nucleation-inducing agent. In fact, we observed the presence of 20 nm-sized calcium phosphate clusters, typical of metastable solutions. Therefore, the sequence of phase transformations from ACP, to OCP until the formation of HAp detected by spectroscopy during *in vitro* MVs mineralization reflects the sequence of phase transitions observed during the precipitation of calcium phosphate from metastable solutions^35^. In line with this interpretation, same previously described factors can extend the lag period of MVs mineralization, such as addition of Mg^2+^ or Zn^2+^ ions^62–64^, also known as inhibitors of HAp crystallization, stabilizing of precursor phases. Interestingly, the addition of ATP can effectively prolong the lag period during the formation and growth of HAp due to its pyrophosphate group, which strongly inhibits mineralization^43^. However, once the ATP is hydrolysed by TNAP, mineralization is enabled to proceed, revealing the primary role of MVs in controlling mineralization by providing a phosphatase machinery.

## 4. Conclusions

The results herein presented combined cryoTEM with bulk characterizations to track the sequence of events that controls MVś driven mineralization in a near-native state. From them, classical experiments and their interpretations were revisited, which enabled us to advance the understanding the mechanisms of MVś action. Our findings revealed that crude MVs isolated from chicken embryos growth plate are heterogeneous and consist not only of bilayered vesicles but also of non-vesicular material, dominated by soluble proteins and fibrillar collagens. The presence of this non-vesicular compounds hinder the precise interpretation of *in vitro* mineralization experiments.

It is important to note that non-vesicular components should not always be considered as contaminants. It has already been reported that the formation of a corona acquired from the extracellular milieu, composed by soluble proteins at the surface of extracellular vesicles can regulate and direct their function^65^. Proteoglycans, collagens, and non-collagenous proteins have been reported to be present in MVs preparations^66^, and the coexistence of MVs with these components within the extracellular matrix, has been long recognized at the ultrastructural level^67^. Therefore, *in vivo* bone mineralization will require interaction among all these components. The results herein presented suggest that the primary role of MVs is to provide an enzymatic machinery to locally control Pi concentration. However, future research should address the individual contribution and interaction between different components in the control of mineralization.

## 5. Methods

### 5.1. Synthetic Cartilage Lymph (SCL) preparation

Synthetic cartilage lymph (SCL) is a solution that mimics the composition of the cartilage fluid^68^. It was prepared by mixing the required salts in ultrapure water (MilliQ^®^) to achieve the following final concentrations: 1.42 mM Na_2_HPO_4_, 1.83 mM NaHCO_3_, 12.7 mM KCl, 0.57 mM MgCl_2_, 100 mM NaCl, 0.57 mM Na_2_SO_4_, 5.55 mM glucose, 63.5 mM sucrose, and 16.5 mM 2-([2-hydroxy-1,1-bis (hydroxymethyl) ethyl]amino)– propanesulfonic acid. The pH of the solution was adjusted to 7.5, at 37°C. SCL was initially prepared without CaCl_2_ and added only just before use to achieve the final concentration required for a given experiment.

### 5.2. Bone tissue digestion and MVs isolation by 3-step differential ultracentrifugation

Leg bones were dissected from chick embryos (17-days post fertilization), characterized to be between developmental stages 42–44 accordingly to Hamburger and Hamilton^69^. In The Netherlands, chick embryos are considered as non-licensed animal use and full animal ethics committee approval was not required for this purpose. The embryos were killed by decapitation, leg bones dissected, and growth plates and epiphyseal cartilages carefully sliced in pieces of approximately 1 mm. The tissue was never frozen and always freshly processed for MVs isolation. MVs were isolated following the protocol described by Buchet et al.^68^ Tissue slices (approximately 4 g) were extensively washed in ice-cold SCL buffer, then digested for 3.5 h, at 37°C, under mild stirring in 16 mL of SCL containing 100 U/mL of collagenase (type II, from *Clostridium histolyticum*, Sigma C6885) and 1 mM CaCl_2_. After digestion, the suspension was filtered through a nylon membrane (100 µm) and consecutively centrifuged at 600 × *g_av_* for 15 min to sediment cells and large debris. The supernatant was then centrifuged at 20 000 x *g_av_* for 20 min using a Ti 70 fixed-angle rotor (Beckman Coulter). Finally, the collected supernatant was centrifuged at 80 000 x *g_av_* for 60 min (Ti 70 fixed-angle rotor). The pellet obtained was suspended in SCL and this preparation we will refer to as crude MVs. Characterizations herein reported were always obtained with freshly isolated MVs that were never subjected to freezing and thawing and stored for no longer than 5 days in ice. We define as a biological replicate the preparation of MVs obtained from bones dissected from 25 animals.

### 5.3. Iodixanol density gradient fractionation of crude MVs

Iodixanol (Optiprep^TM^ 60 vol.% in water, Serumwerk Bernburg, Germany) was prepared diluted in ice-cold SCL to achieve a final concentration 48 vol.%. This working-solution was then mixed with a dispersion of crude MVs in SCL to achieve a final concentration of 45 vol.%. 4 mL of the iodixanol solution containing the crude MVs was loaded in the bottom of an ultra-clear tube and then solutions of descending densities (35, 30, 28, 26, 24, 22, 20 and 10 %) were carefully layered on top. All solutions with different densities were prepared in SCL containing 1 mM CaCl_2_, to prevent dissolution of any labile Ca^2+^ and Pi initially present^49^. The bottom-loaded 45-10% gradient was then centrifuged at 120 000 x *g_av_* for 16 h in a SW40 Ti Swinging Bucket rotor (Beckman Coulter). After ultracentrifugation, fractions were carefully collected from the top of the gradient, and 12-fold diluted in ice-cold SCL and subjected to ultracentrifugation at 100 000 x *g_av_* for 2 h in a Ti 70 fixed-angle rotor (Beckman Coulter). The resulting pellets were resuspended in SCL and freshly used for downstream analysis and never subjected to freezing and thawing.

### 5.4. Protein quantification and alkaline phosphatase activity

Total protein concentration was quantified using the Bradford protein assay (Biorad), using bovine serum albumin as standard. Alkaline phosphatase activity was determined by the hydrolysis of p-nitrophenyl phosphate (p-NPP, Sigma-Aldrich) in a reaction medium containing 10 mM p-NPP and 1 mM MgCl_2_ in AMPOL buffer (2-amino-2-methyl-1-propanol), pH 10.3, at 37°C. The reaction was monitored by changes in the absorbance at 405 nm related to the formation of the yellowish product p-nitrophenolate (p-NP^-^) and the specific activity expressed as µmol pNP^−^/min/mg of total protein.

### 5.5. SDS-PAGE and western blotting

Samples were prepared in Laemmli buffer and heated at 95°C for 10 min before being loaded on gels^70^. The samples were separated on 4%–15% Mini-PROTEAN® TGX™ Precast Gels (Biorad) under reducing conditions before being transferred to PVDF membranes (Biorad) using a Trans-Blot® Turbo Blotting System. All procedures were done using manufacturers protocols. Membranes were blocked for 1 h in Odyssey TBS Blocking Buffer (LI-COR Biosciences, Lincoln, NE, USA), and incubated with anti-ALPL primary antibody (Proteintech, 11187-1-AP), diluted 1:1000, overnight. For fluorescence detection of proteins, IRDye 800CW anti-rabbit IgG (H+L) secondary antibody (LI-COR) was used. Detection was with an Odyssey Imaging System (LI-COR).

### 5.6. Nanoparticle tracking analysis (NTA)

MVs size distribution and concentration were estimated in a NanoSight NS300 equipped with a syringe pump (Malvern Panalytics, Malvern, UK). Concentrations were calculated using Nanoparticle Tracking Analysis 3.2 software (Nanosight Ltd, Amesbury, UK). Vesicles were diluted in PBS until a suitable concentration for analysis was reached (20-60 particles per frame). Each sample was measured for 30 s, using the following software settings: flow rate 50, camera level 10, and detection threshold 5.

### 5.7. CryoTEM sample preparation and imaging

3 μL of sample was applied on glow-discharged Au200 R2/1 grids (Quantifoil), and the excess liquid was removed by blotting for 4 s (blot force 3) using filter paper followed by plunge freezing in liquid ethane using a FEI Vitrobot Mark IV, at 100% humidity and 25°C. A JEOL JEM-2100 microscope (Jeol Ltd., Tokyo, Japan) equipped with a Gatan 914 high tilt cryo holder and a LaB6 filament was used for cryogenic imaging at 200 kV. Images were recorded with a Gatan 833 Orius camera (Pleasanton, CA, USA).

### 5.8. Mineralization experiments

Crude and low-dense MVs were exposed to SCL medium supplemented with CaCl_2_ (2 mM) and either 2 mM Na_2_HPO_4_ or 1 mM adenosine triphosphate (Sigma-Aldrich). Mineralization was carried out at 37°C and using samples with fixed 50 µg of total protein/mL concentration. Turbidity was measured as absorbance at 350 nm, in 200 µL of solution in 96-well plate using a Biorad model 680 microplate reader. Dynamic light scattering (DLS) was carried out in a NanoZS equipament (Malvern, UK).

For on-grid mineralization experiments, 10 µL of samples were placed on top of a glow-discharged Au200 R2/1 grids (Quantifoil) and incubated in a 100%-humidity controlled environment, at 37°C. After incubation for a desired time, samples were processed for vitrification and imaging as described above.

## Acknowledgments

M.A.E.C, P.C. and A.P.R. are supported by The São Paulo Research Foundation (FAPESP) (grants 2019/26059-8, 2019/25054-2 and 2019/08568-2). N.S. is supported by an European Research Council (ERC) Advanced Investigator grant (H2020-ERC-2017-ADV-788982-COLMIN). A.A. was in part supported by a VENI grant from the Netherlands Scientific Organization NWO (VI.Veni.192.094).

