## Supplementary Figures for "Unravelling the relationship between isolated bone matrix vesicles and forming mineral at the nanometer scale"

**Figure S1. TEM of crude and low-dense MVs.** Samples were fixed with 1% glutaraldehyde, dropped in a TEM grid and stained with uranyl acetate 1%. a) crude MVs were we can detect the presence of fibrillar proteins. b) Typical cup-shaped morphology of dried vesicles. c) Low-dense MVs, showing the removal of associated fibrillar proteins. Notice that due to staining is difficult to discern their content, i.e. if there is any electron-dense mineral associated with the vesicles. Scale bars, 200 nm

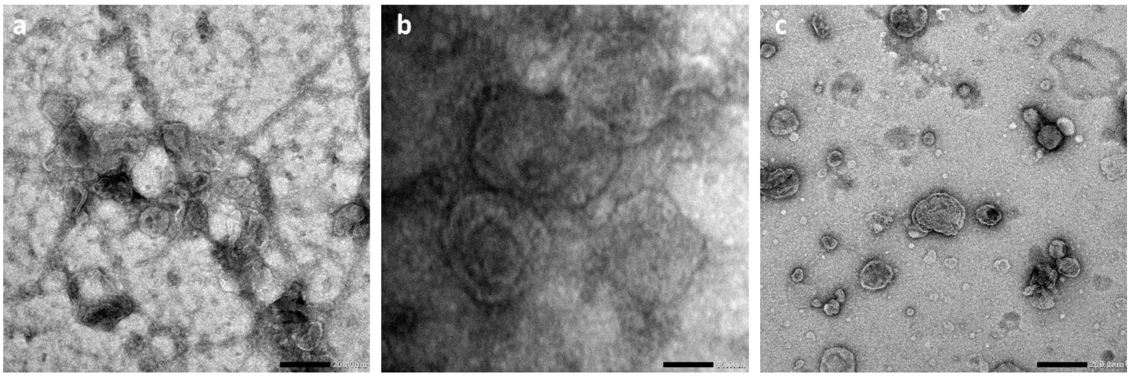

**Figure S2. Mineralization triggered by non-vesicular particles**. High-density fraction trigger mineral formation, observed after 24 h, when incubated at same conditions where low-dense MVs did not. Scale bars, 100 nm

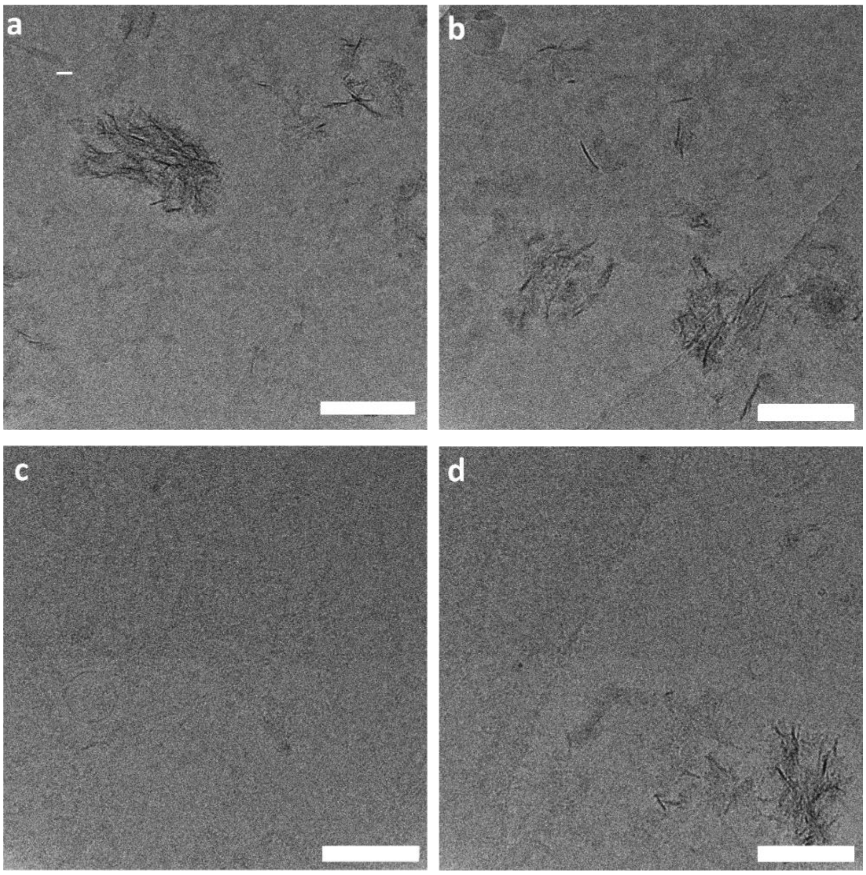
